# Design of Epitope Based Peptide Vaccine against Brucella Abortus OmpW Family Protein using Immunoinformatics

**DOI:** 10.1101/2021.12.27.474190

**Authors:** Mustafa Elhag, Abdelrahman Hamza Abdelmoneim, Anfal Osama Sati, Moaaz Mohammed Saadaldin, Nagla Mohammad Ahmad, Mohammed A. Hassan

## Abstract

*Brucella abortus* is a small aerobic, non-spore-forming, non-motile intracellular *coccobacilli* localized in the reproductive organs of host animals and causes acute or chronic disorders. It infects approximately 200 cases per 100,000 of the population and has become endemic in many countries. OmpW family protein is an outer membrane protein involved in the initial interaction between the pathogen and it’s host. This study predicts an effective epitope-based vaccine against OmpW family protein of *Brucella abortus* using immunoinformatics tools. Sequences were obtained from NCBI and prediction tests were accomplished to analyze possible epitopes for B and T cells. Seven B cell epitopes passed the antigenicity, accessibility and hydrophilicity tests. Forty-three MHC I epitopes were the most promising, while 438 from MHC II. For the population coverage, the epitopes covered 99.97% of the alleles worldwide excluding certain MHC II alleles. We recommend *invivo* and *invitro* studies to prove it’s effectiveness.

## 1. Introduction

Brucellosis is a zoonotic infection caused by the bacterial genus *Brucella*. The disease is an old one and has been known by various names, including Mediterranean fever, Malta fever, gastric remittent fever, and undulant fever. Humans are the accidental hosts. In humans, they are characterized by a variable incubation period (ranging from several days up to several months), and clinical signs and symptoms of continued, intermittent or irregular fever of variable duration with headaches, weakness, profuse sweating, chills, depression and weight loss. Localized suppurative infections may also occur. The course of the disease can be variable, especially in persons either not or inadequately treated. Diagnosis of clinical brucellosis in humans and animals is initially made by use of appropriate serological or other immunological tests, and confirmed by bacteriological isolation and identification of the agent, but brucellosis continues to be a major public health concern worldwide and is the most common zoonotic infection [1-3].

*Brucella* organisms, which are small aerobic intracellular *coccobacilli*, localize in the reproductive organs of host animals, causing acute or chronic disorder. They are spread widely in the animal’s urine, milk, placental fluid and other fluids. Till date, eight species have been identified, named primarily for the source animal or features of infection. Of these, four moderate-to-significant human pathogenicity species are *Brucella melitensis* (from sheep; highest pathogenicity), *Brucella suis* (from pigs; high pathogenicity), *Brucella abortus* (from cattle; moderate pathogenicity), and *Brucella canis* (from dogs; moderate pathogenicity) [3].

*B. abortus* is a gram-negative alpha-proteobacterium in the family *Brucellaceae* and is one of the causative agents of brucellosis. The rod-shaped pathogen, classified under the domain prokaryotic bacteria, is non-spore-forming, non-motile and aerobic. The mortality in recognized symptomatic acute or chronic cases of brucellosis is very low, certainly less than 5% and probably less than 2%. *B. abortus* primarily affects cattle, other biovidae and cervidae. It is usually the result of the rare instance of *Brucella* endocarditis or is the result of severe CNS involvement, often as a complication of endocarditis. Post-mortem analysis confirms that the burden of acute brucellotic infection is borne by tissues of the lymphoreticular system. Members of the genus can grow on enriched media like blood agar or chocolate agar. Identification in species level can be done by agglutination with monospecific serum, cultivating the strains in the presence of dyes and/or with PCR methods. They are catalase, oxidase and urea positive bacteria [1-4]. It is an intracellular coccobacilli pathogen that infects genitourinary tract of cattle and humans who are infected after exposure to *B. abortus* or was previously identified to be immunogenic in animals infected with *Brucella species* to infected animals or contaminated meat or dairy products. The response against *B. abortus* involves the whole gamut of the immune system, from innate to adaptive immunity resulting from stimulation of antigen-presenting cells, NK cells, CD4+ and CD8+ T cells, and B cells [5].

Brucellosis is an important human disease in many parts of the world, as much as 200 cases per 100,000 of the population in some regions of the world; besides, the infection has become endemic in many countries [6]. It is a bacterial infection which is known as undulant fever, Mediterranean fever, or Malta fever. It is a zoonosis and the infection is almost invariably transmitted to people by direct or indirect contact with the infected animal (eg eating raw or unpasteurized dairy products or air) [7].

To treat or cure this infection, we need to identify its outer membrane proteins such as OmpW family protein. However, most of the gram negative bacteria have evolved many mechanisms of attaching and invading host epithelial and immune cells. In particular, many outer membrane proteins (OMPs) are involved in this initial interaction between the pathogen and it’s host. This can make a useful use of it in the designing vaccine. A number of small pore-forming OMPs are all composed of eight stranded β-barrel proteins and include members of the OmpA, OmpW with a beta-barrel structure consisting of eight non-parallel beta strands. OmpW family are widely distributed among gram-negative bacteria. These proteins, together with the related OmpA-like peptidoglycan associated lipoproteins, are involved in interactions with host cells and are mediators of virulence. In many cases, these proteins interact with host immune cells and can be considered as pathogen associated molecular patterns (PAMPS) due to their ability to signal through toll like receptor molecules and other pattern recognition receptors [8]. In some studies, they discovered that the 14-kDa protein possessed immunoglobulin binding and hemagglutination properties that appeared to be based on the protein’s lectin-like properties. Hemagglutination inhibition experiments suggested that the 14-kDa protein has affinity towards mannose. Disruption of the gene encoding the 14-kDa protein in virulent *B. abortus* strain 2308 induced a rough-like phenotype with an altered smooth lipopolysaccharide (LPS) immunoblot profile and a significant reduction in the bacterium’s ability to replicate in mouse spleens. However, the mutant strain was stably maintained in mouse spleens at 2.0 to 2.6 log(10) CFU/spleen from day 1 to week 6 after intra-peritoneal inoculation with 4.65 log(10) CFU. In contrast to the case for the smooth virulent strain 2308, in the rough attenuated strain RB51 disruption of the 14-kDa protein’s gene had no effect on the mouse clearance pattern. These findings indicate that the 14-kDa protein of *B. abortus* possesses lectin-like properties and is essential for the virulence of the species, probably because of its direct or indirect role in the synthesis of smooth LPS [9]. The vaccine strains are the most commonly used to protect livestock against infection and abortion. However, due to some disadvantages of these vaccines, numerous studies have been conducted for the development of effective vaccines that could also be used in other susceptible animals. In this article, we compared different aspects of immunogenic antigens that have been a candidate for the brucellosis vaccine to get the most effective, efficient, active and harmless one [10].

## 2. Materials and methods

### Protein Sequence Retrieval

The sequences of the *B. abortus* strains were retrieved from the National Center for Biotechnology Information (NCBI) database in May 2019 in FASTA format. These strains were collected from different parts of the world for immunoinformatics analysis. The retrieved protein strains had a length of 227 with the name OmpW family protein.

### Conserved region identification

BioEdit Sequence Alignment Editor Software (version 7.0.5.3) was used to determine the conserved regions of the retrieved sequences of *B. abortus* OmpW family using Clustal-W multiple sequence alignment (MSA). Amino acid composition and Molecular weight of the protein were also obtained.

### B cell Epitope Prediction

The prediction of Linear B cell epitopes is done by objected reference sequence of *B. abortus* to Bepipred Linear Epitope Prediction tool 2.0 at Immune Epitope Database and Analysis Resource IEDB (http://tools.iedb.org/bcell/) [11]. Bioedit sequence alignment editor was used to identify the epitope Conservancy. Only epitopes with 100% conservancy were selected and analysed for surface antigenicity through Kolaskar & Tongaonkar Antigenicity tool, Emini Surface Accessibility Prediction tool for surface accessibility and Parker Hydrophilicity Prediction tool for hydrophilic, accessible, or mobile regions with thresholds of 1.033, 1.000 and 0.779 respectively [12-14]. Epitopes that pass all tests were predicted as B cell epitope.

### T cell Epitope Prediction MHC Class I Binding

Analysis of peptide binding to the MHC (Major Histocompatibility complex) class I molecule was calculated by the IEDB MHC I prediction tool (http://tools.iedb.org/mhci/) to predict cytotoxic T cell epitopes. To predict the binding affinity, we used Artificial Neural Network (ANN) 4.0 prediction method [15, 16]. Before the binding analysis, all the alleles were selected, and the length was set to 9 amino acids before the prediction was done. Then, the conserved epitopes were classified according to their half-maximal inhibitory concentration (IC50) into high affinity (IC50<100), moderate affinity (IC50<500) and low-affinity epitopes (IC50 <500). Only high-affinity epitopes with their corresponding alleles were subjected to population coverage analysis.

### T cell Epitope Prediction MHC Class II Binding

Prediction of T cell epitopes was identified by using the IEDB MHC II prediction tool (http://tools.iedb.org/mhcii/) consuming the NN-align method [17]. To determine the interaction potentials of T cell epitopes and their respective MHC class II alleles, all human allele references were set. All conserved epitopes that bind to alleles at a score less than 500 half-maximal inhibitory concentration (IC50<500) were selected for further analysis.

### Prediction of Allergens

AllerTop 2.0 (http://www.ddg-pharmfac.net/AllerTOP/), an online server, was used to analyze the predicted allergenicity of the selected epitope based on the main physiochemical properties of proteins. The predicted epitopes were classified as either “Probable Allergen” or “Probable Non-allergen” [18].

### Population Coverage

Population coverage analysis tool of IEDB was used for predicting the population coverage of the whole world (http://tools.iedb.org/tools/population/iedb_input) [19]. Given a set of epitopes with their corresponding HLA alleles, this tool calculates the fraction of individuals predicted to cover. Epitopes with the highest frequency were selected for modelling.

### Homology Modelling

RaptorX (https://www.raptor.uchicago.edu) was used to predict the 3D structure of *B. abortus* OmpW family protein by comparing it to a set of homologs. For visualization and analysis of the molecular structure of the obtained 3D protein structure, UCSF Chimera (version1.8) (http://www.cgl.ucsf.edu/chimera) was used.

## 3. Results

### Multiple sequence alignment

Sixty-one sequences of the OmpW family protein of *B. abortus* were subjected to multiple sequence alignment against the reference protein to look for the conserved regions.

### B-cell epitope prediction

Forty-seven epitopes passed the four B-cell prediction tools (Bepipred linear epitope 2, Emini surface accessibility (threshold of 1), Kolaskar & Tongaonkar antigenicity (threshold of 1.033), Parker hydrophilicity prediction (threshold of 0.779)) and the Allertop test. Below is the result of the top seven B-cell epitopes that had passed all four tests and have the largest length (Table 1).

**Table 1:**
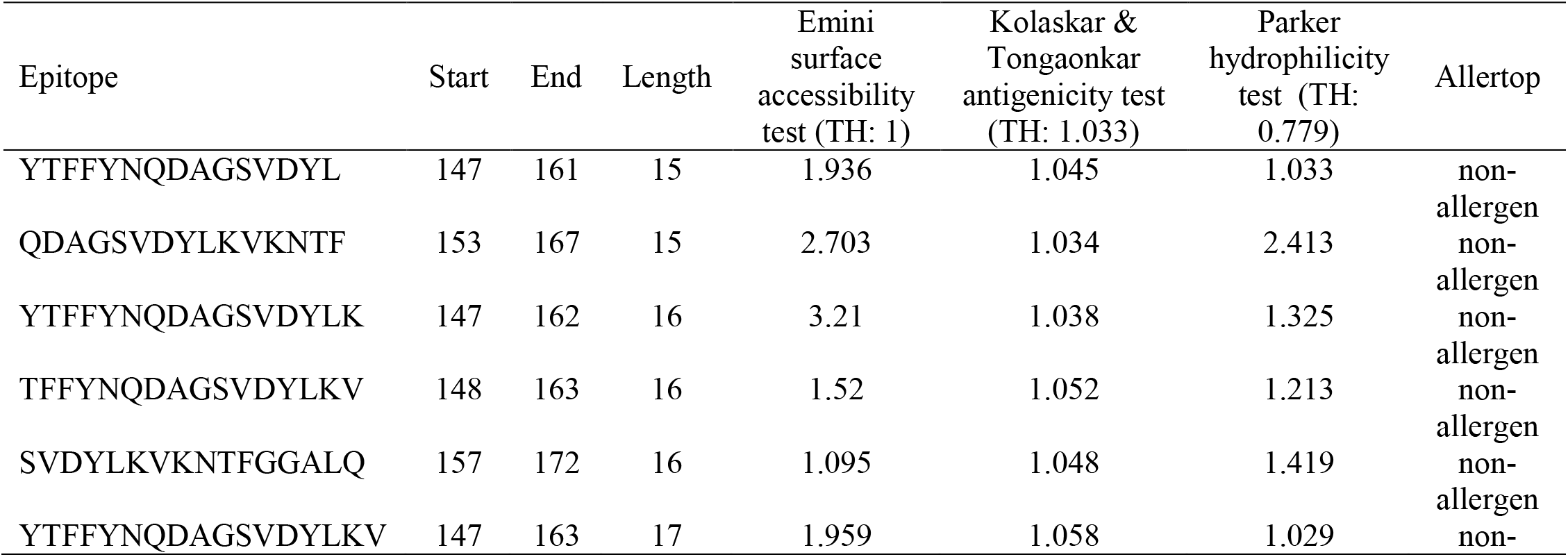
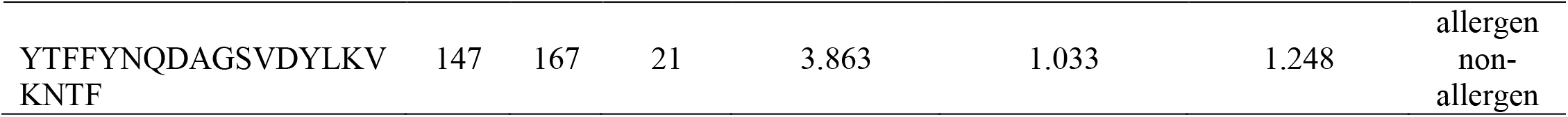
Illustrates the largest seven B-cell epitopes that passed all immunogenicity tests.

### T- Cell epitope prediction: MHC class-I binding peptides

The reference sequence was analyzed using (IEDB) MHC class I binding prediction tool to predict for possible T cell epitopes interacting with different MHC class I alleles with IC50 <100. Forty-three peptides were predicted to interact with different MHC class I alleles. All conserved epitopes with their corresponding MHC class I alleles and IC50 scores are shown in (Table 2).

**Table 2:**
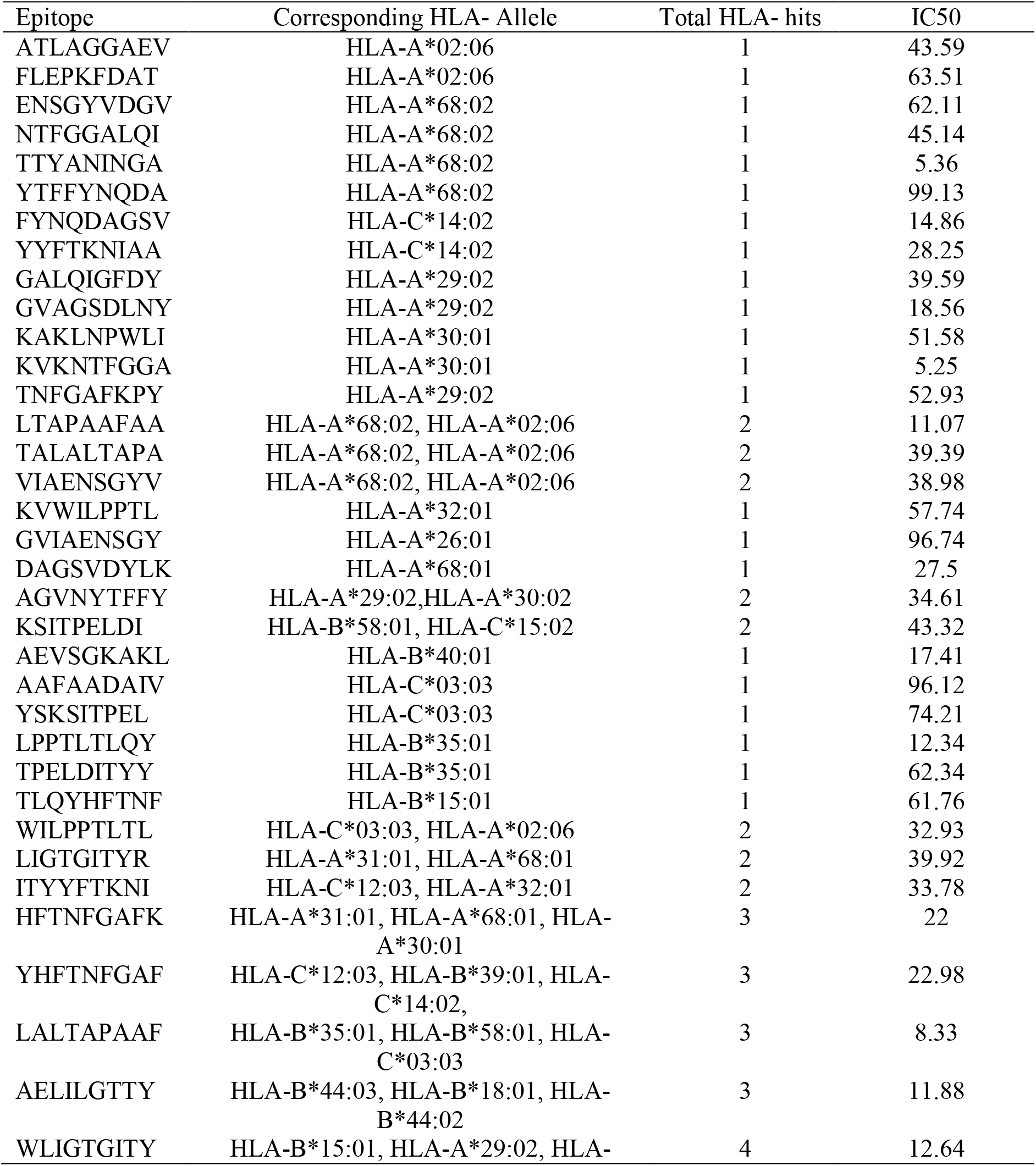
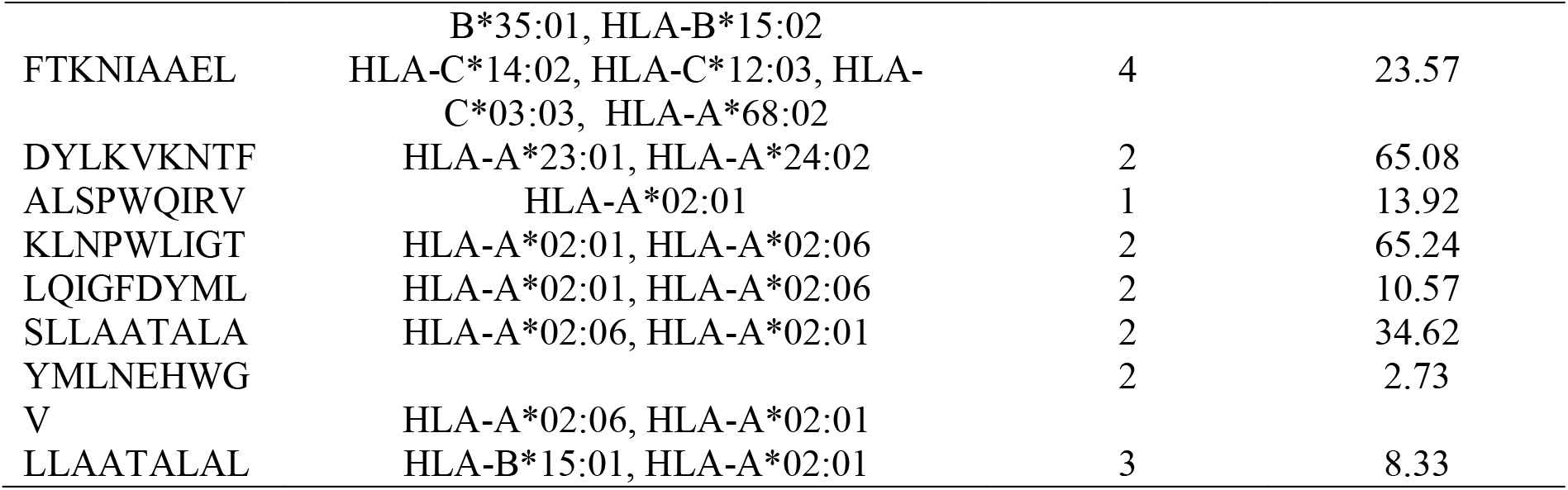
Shows all conserved T- Cell peptides which interact with MHC class I and their corresponding IC50 scores.

### T- Cell epitope prediction: MHC class-II binding peptides

The reference sequence was analyzed using (IEDB) MHC class II binding prediction tool. There were 438 predicted epitopes found to interact with MHC class II alleles. The most promising epitopes with the number of HLA hits and their IC50 scores were shown in (Table 3).

**Table 3:** The most promising T- Cell epitopes interacting with MHC-II alleles.

| Epitope | No. of HLA- hits | IC50 score |
| --- | --- | --- |
| GVNYTFFYNQDAGSV | 18 | 16.2 |
| IAAELILGTTYANIN | 18 | 17.1 |
| NIAAELILGTTYANI | 18 | 17.7 |
| AELILGTTYANINGA | 18 | 23.2 |
| LQIGFDYMLNEHWGV | 18 | 24 |
| VNYTFFYNQDAGSVD | 19 | 16.9 |
| ALQIGFDYMLNEHWG | 19 | 29.2 |
| GALQIGFDYMLNEHW | 20 | 28.6 |
| ELDITYYFTKNIAAE | 21 | 8.6 |
| PELDITYYFTKNIAA | 21 | 10.5 |
| YFTKNIAAELILGTT | 21 | 16.7 |
| LLAATALALTAPAAF | 22 | 11.1 |
| SLLAATALALTAPAA | 22 | 11.5 |
| KSLLAATALALTAPA | 23 | 12 |
| NRFTKSLLAATALAL | 24 | 6.9 |
| MNRFTKSLLAATALA | 24 | 7.6 |
| FTKSLLAATALALTA | 24 | 10.7 |
| TKSLLAATALALTAP | 24 | 13.3 |
| RFTKSLLAATALALT | 25 | 7.7 |
| YYFTKNIAAELILGT | 26 | 6.3 |
| DITYYFTKNIAAELI | 27 | 4.6 |
| TYYFTKNIAAELILG | 27 | 5.3 |
| LDITYYFTKNIAAEL | 27 | 5.5 |
| ITYYFTKNIAAELIL | 30 | 4.7 |

### Population coverage analysis

All MHC class I and II epitopes were assessed for global population coverage using IEDB population coverage tool. For MHC I, the epitope with the highest global population coverage was LLAATALAL with coverage percentage of 45.62% (Table 4 and Figure 2).

**Table 4:** The most promising MHC I binding peptides with the highest global population coverage percentages

| Epitopes | Coverage |
| --- | --- |
| LLAATALAL | 45.62% |
| KLNPWLIGT | 40.60% |
| LQIGFDYML | 40.60% |
| SLLAATALA | 40.60% |
| YMLNEHWGV | 40.60% |
| Total global coverage | 97.4% |

MHC II epitope with the highest global population coverage was ITYYFTKNIAAELIL with a 99.97% coverage (Table 5 and Figure 3).

**Table 5:** Showing most promising MHC II binding epitopes with the highest global population coverage

| Peptide | Coverage |
| --- | --- |
| ITYYFTKNIAAELIL | 99.97% |
| DITYYFTKNIAAELI | 99.94% |
| TYYFTKNIAAELILG | 99.94% |
| YYFTKNIAAELILGT | 99.92% |
| TKSLLAATALALTAP | 99.90% |
| LDITYYFTKNIAAEL | 99.90% |
| NRFTKSLLAATALAL | 99.86% |
| RFTKSLLAATALALT | 99.86% |
| KSLLAATALALTAPA | 99.85% |
| FTKSLLAATALALTA | 99.85% |
| SLLAATALALTAPAA | 99.80% |
| MNRFTKSLLAATALA | 99.80% |
| YFTKNIAAELILGTT | 99.69% |
| PELDITYYFTKNIAA | 99.64% |
| ELDITYYFTKNIAAE | 99.53% |
| IAAELILGTTYANIN | 99.51% |
| NIAAELILGTTYANI | 99.51% |
| AELILGTTYANINGA | 99.51% |
| GALQIGFDYMLNEHW | 99.48% |
| ALQIGFDYMLNEHWG | 99.27% |
| Total global coverage | 99.99% |

### Homology modelling

Raptor X homology modeler software used for prediction of the most identical OmpW protein structure of *B. abortus*, and the PDB ID obtained was 2f1tA. The most promising peptides 3D structure was afterwards, edited and visualized using the Chimera software (version 1.13.1rc).

## 4. Discussion

In the current study, we proposed different peptides that can be recognized by B and T cells to produce vaccine against OmpW family protein of *Brucella abortus*. Immunoinformatics based peptide vaccines overcome the side effects of conventional vaccines. Peptide vaccines require less time and cost to be produced, can stimulate effective immune response and can target rapidly mutating pathogens [20, 21].

The reference sequence of OmpW family protein of *B. abortus* was subjected to Bepipred linear epitope prediction 2 test, Emini surface accessibility test, Kolaskar and Tongaonkar antigenicity test, Parker hydrophilicity test and Allertop to determine the binding ability to the B cell and to test the surface accessibility, immunogenicity, hydrophilicity and allergenicity respectively. Furthermore, multiple sequence alignment was used to study the conservation of the protein against sixty similar protein sequences which showed a high degree of conservation except at three sites (29, 51 and 192), which signify the necessity of this protein for the survival of *B. abortus* species (Figure 1). Forty-seven peptides passed four B-cell prediction tools (Table 1). Bepipred linear epitope 2, Emini surface accessibility, Kolaskar & Tongaonkar antigenicity, Parker hydrophilicity prediction, and Allertop methods. Of these peptide seven were promising as they passed all prediction tools. These peptide has the ability to produce effective antibody response.

**Figure 1:**
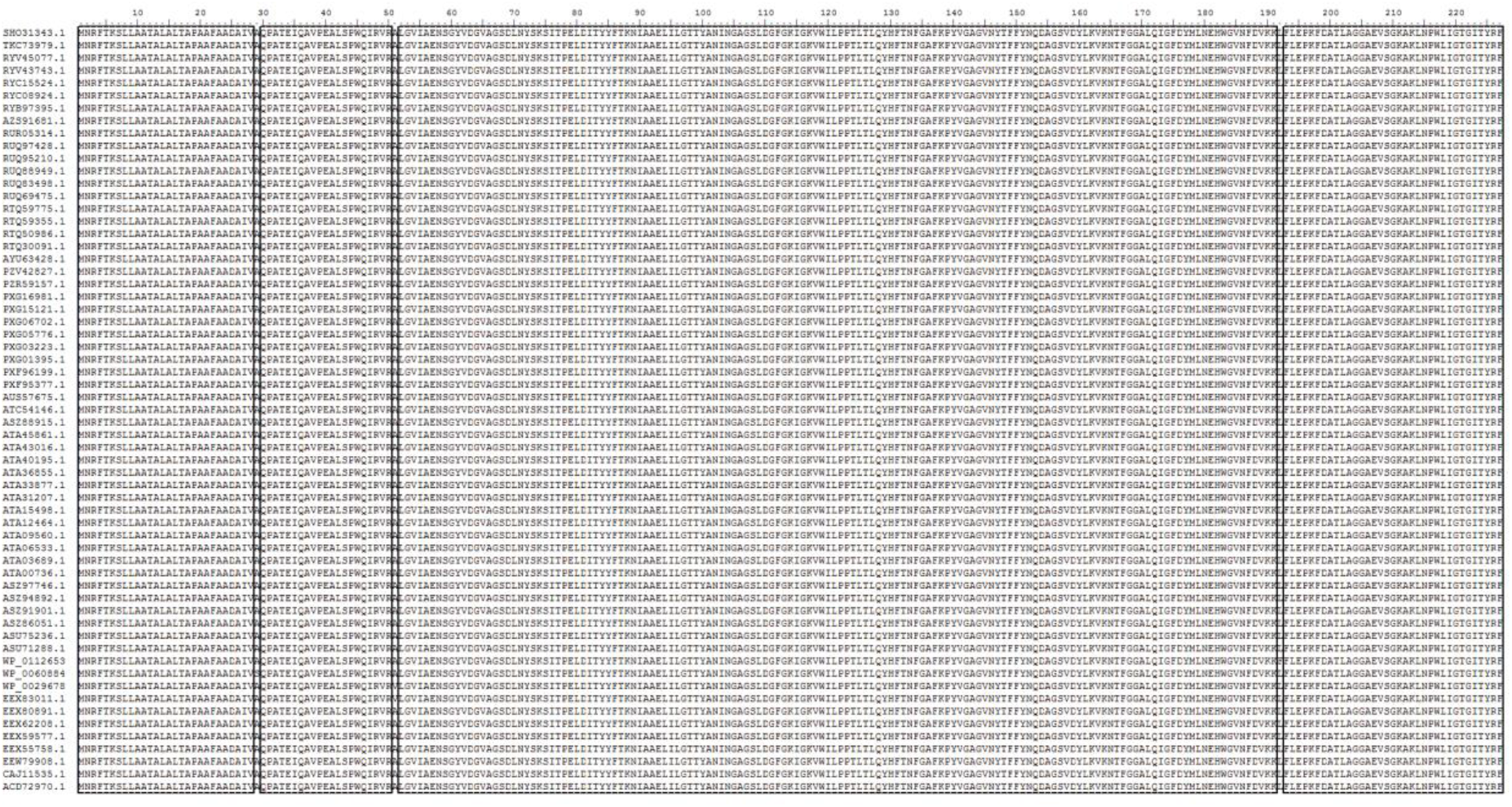
Illustrates the multiple sequence alignment of sixty-one proteins of OmpW family protein of *B. abortus*, showing highly conserved regions in the sequence except in positions 29, 51 and 192 (using bioedit software).

**Figure 2:**
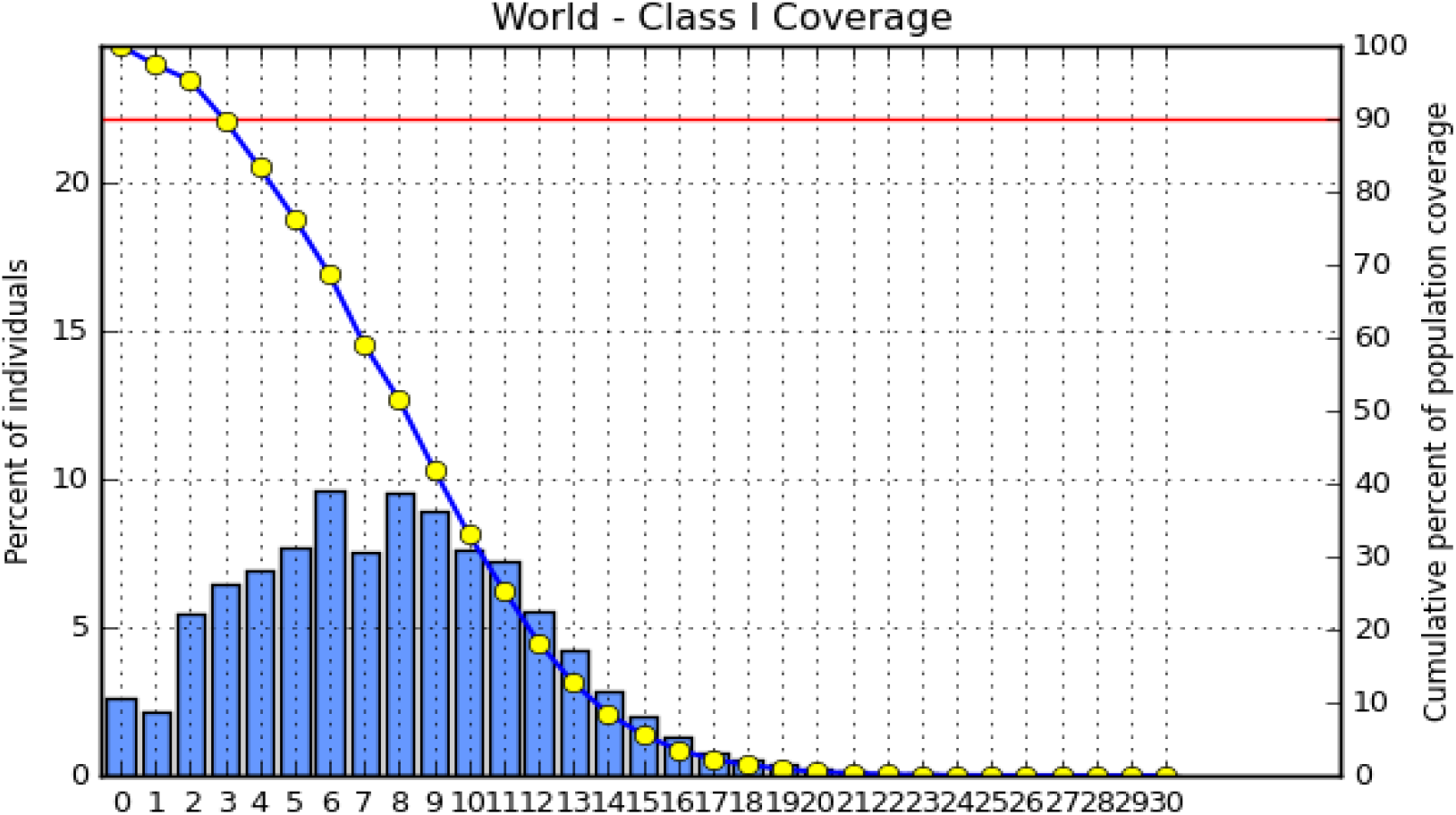
Illustrates global population coverage of T- Cell epitopes of OmpW protein of *B. abortus* which interacts with MHC I alleles

**Figure 3:**
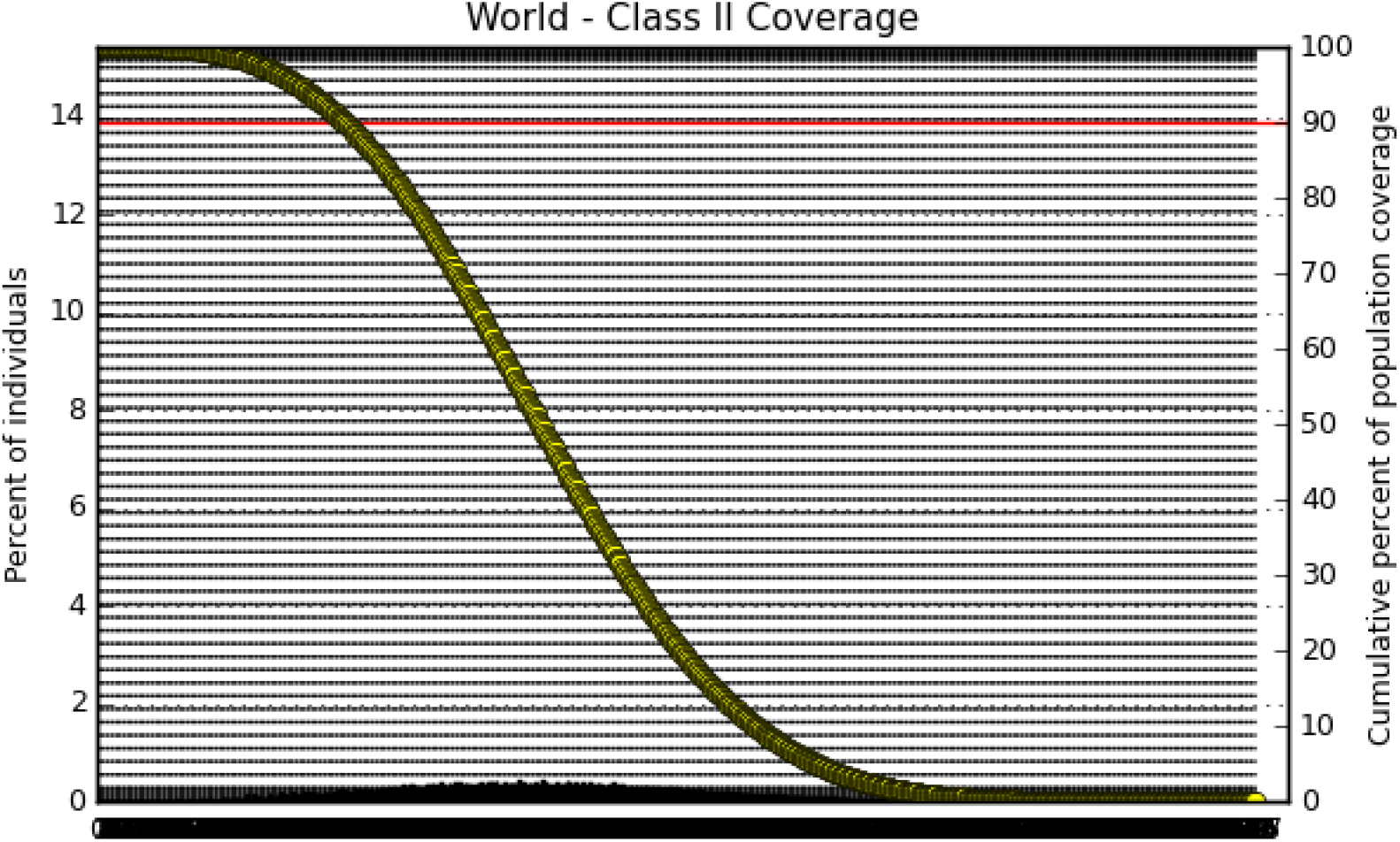
Illustrates global population coverage of T- Cell peptides of OmpW protein of *B. abortus* which interacts with MHC II alleles

The reference sequence was further analyzed using IEDB MHC I and II binding prediction tools to predict T cell epitopes. Forty-three peptides were predicted to interact with MHC I alleles with IC50 <=100 (Table 2). Seven of them were most promising and had the ability to bind to the highest number of MHC I alleles (HFTNFGAFK, YHFTNFGAF, LALTAPAAF, AELILGTTY, WLIGTGITY, FTKNIAAEL and LLAATALAL). Forty three predicted epitopes interacted with MHC II alleles with IC50 <= 500. Twenty four of them were most promising and had the ability to bind to the highest number of MHC II alleles (Table 3).

The best epitope with the highest population coverage for MHC class I was LLAATALAL with 45.62% in two HLA hits, one of them is HLA-A*02:01 - the worldwide predominant MHC I allele which is capable of inducing powerful CTL responses and the coverage of population set was 97.4% for the whole MHC I epitopes (Table 4). Regarding population coverage for MHC class II, the best epitope was ITYYFTKNIAAELIL scoring 99.97% with thirty HLA hits and the coverage of population set was 99.99% for the whole MHC II epitopes (Table 5). This high coverage percentage makes these peptides excellent targets for the vaccine design. The combined coverage for both MHC I and MHC II is 100%, which further solidify the significance of these epitopes (Figure 4).

**Figure 4:**
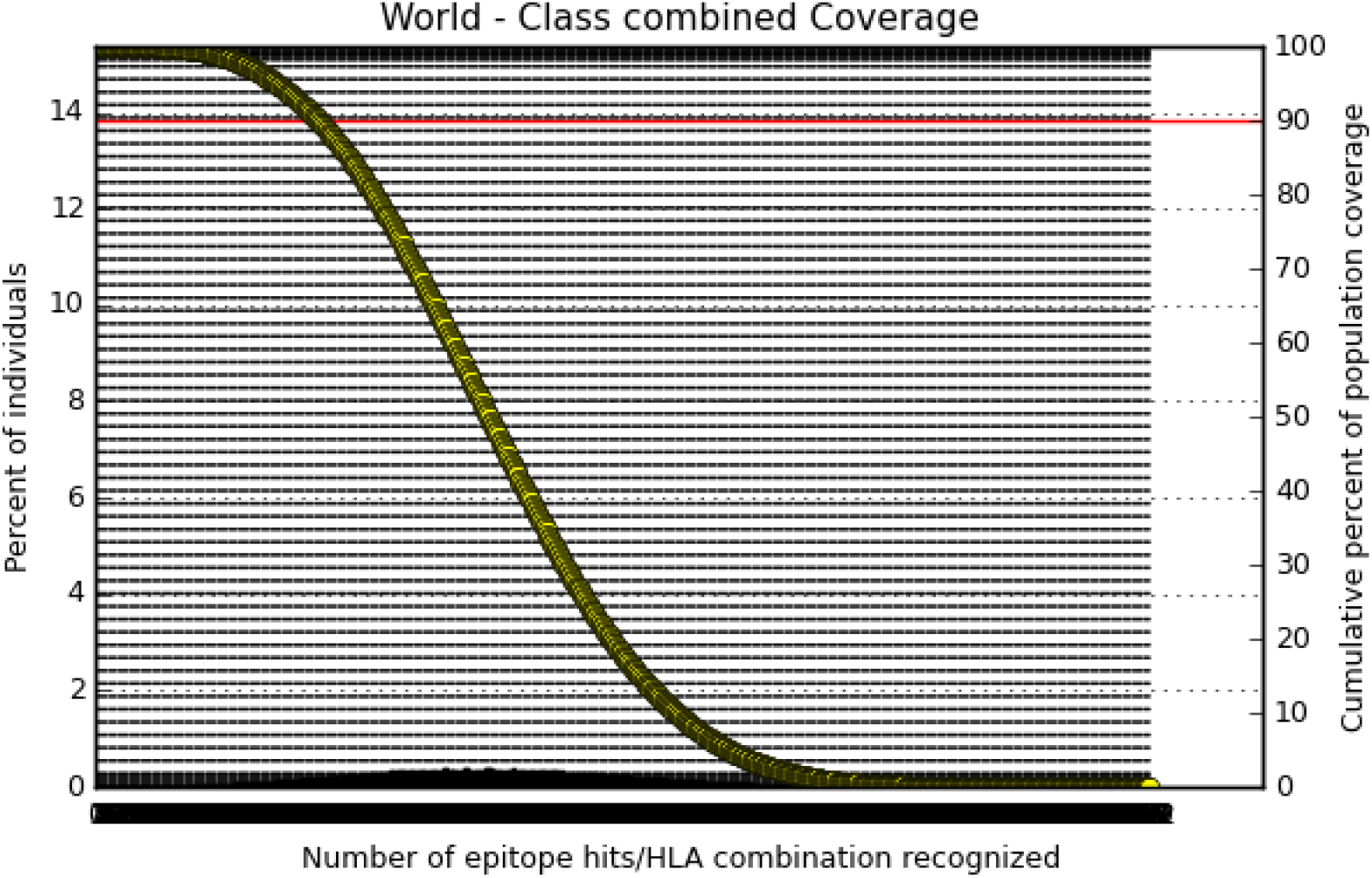
Illustrates global population coverage of T- Cell peptides of OmpW protein of *B. abortus* which interacts with both MHC class I and II alleles

**Figure 5:**
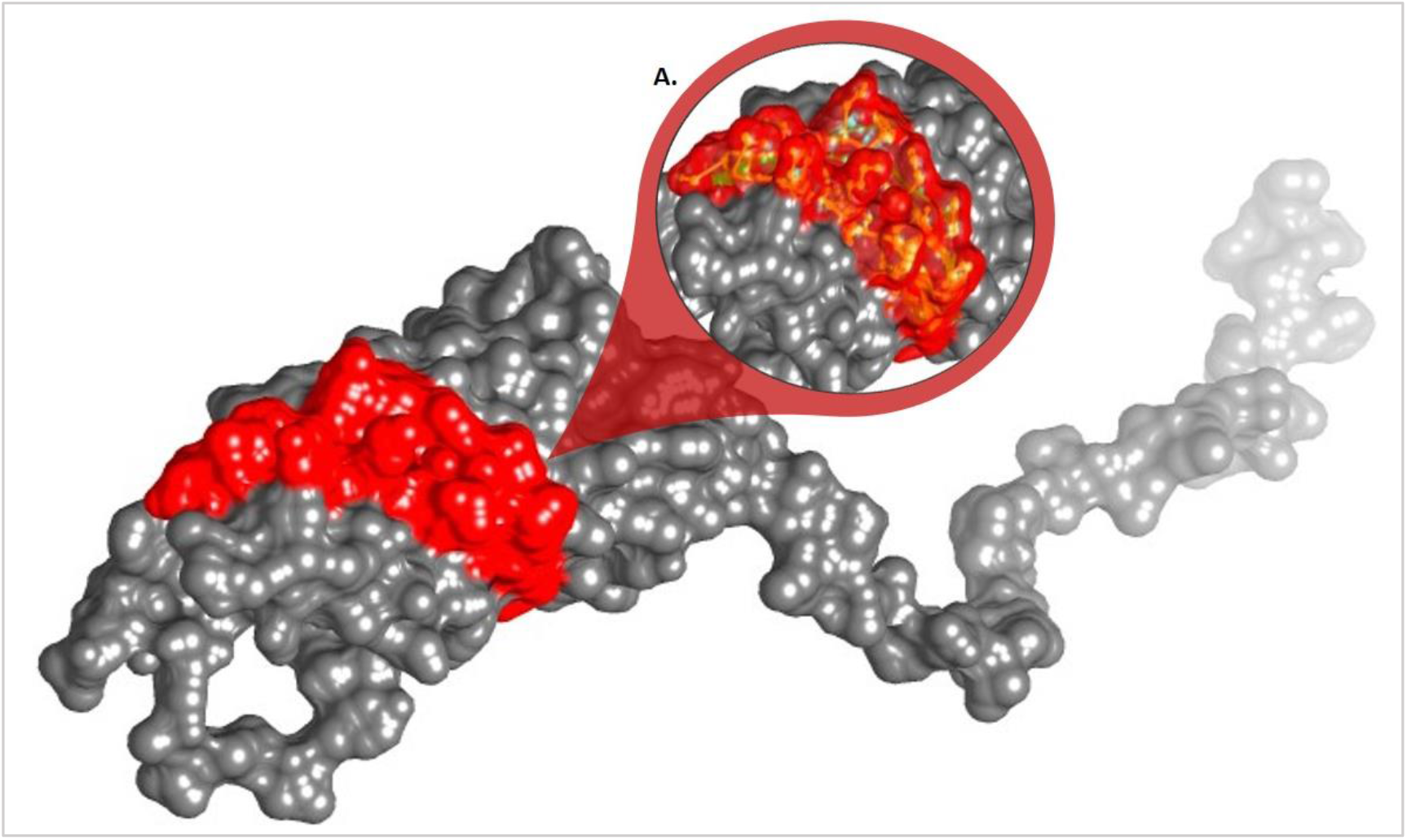
Three-Dimensional structure of OmpW protein of *B. abortus* showing most promising B-cell peptides located in the same conserved area. Starting from position 147 to position 172 (using chimera 1.13.1rc)

**Figure 6:**
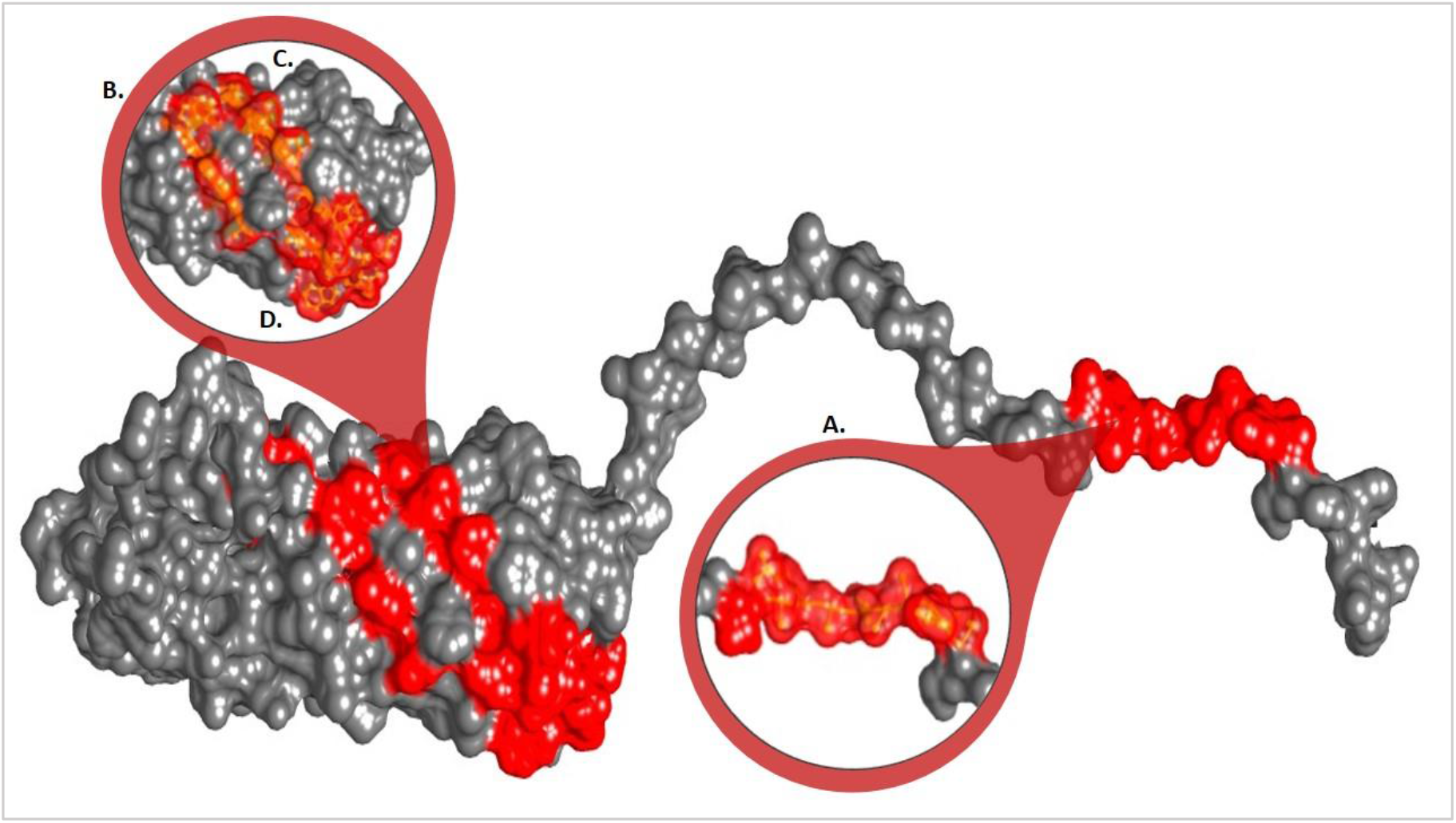
Three-Dimensional structure of OmpW protein of *B. abortus* showing most promising T- Cell peptides interacting with MHC-I alleles, and located in the same conserved area. A. shows peptides LLAATALAL and SLLAATALA (positions from 6 to 15). B. KLNPWLIGT (from 213 to 221). C. LQIGFDYML (from 171 to 179) and D. YMLNEHWGV (from 177 to 185) (using chimera 1.13.1rc)

**Figure 7:**
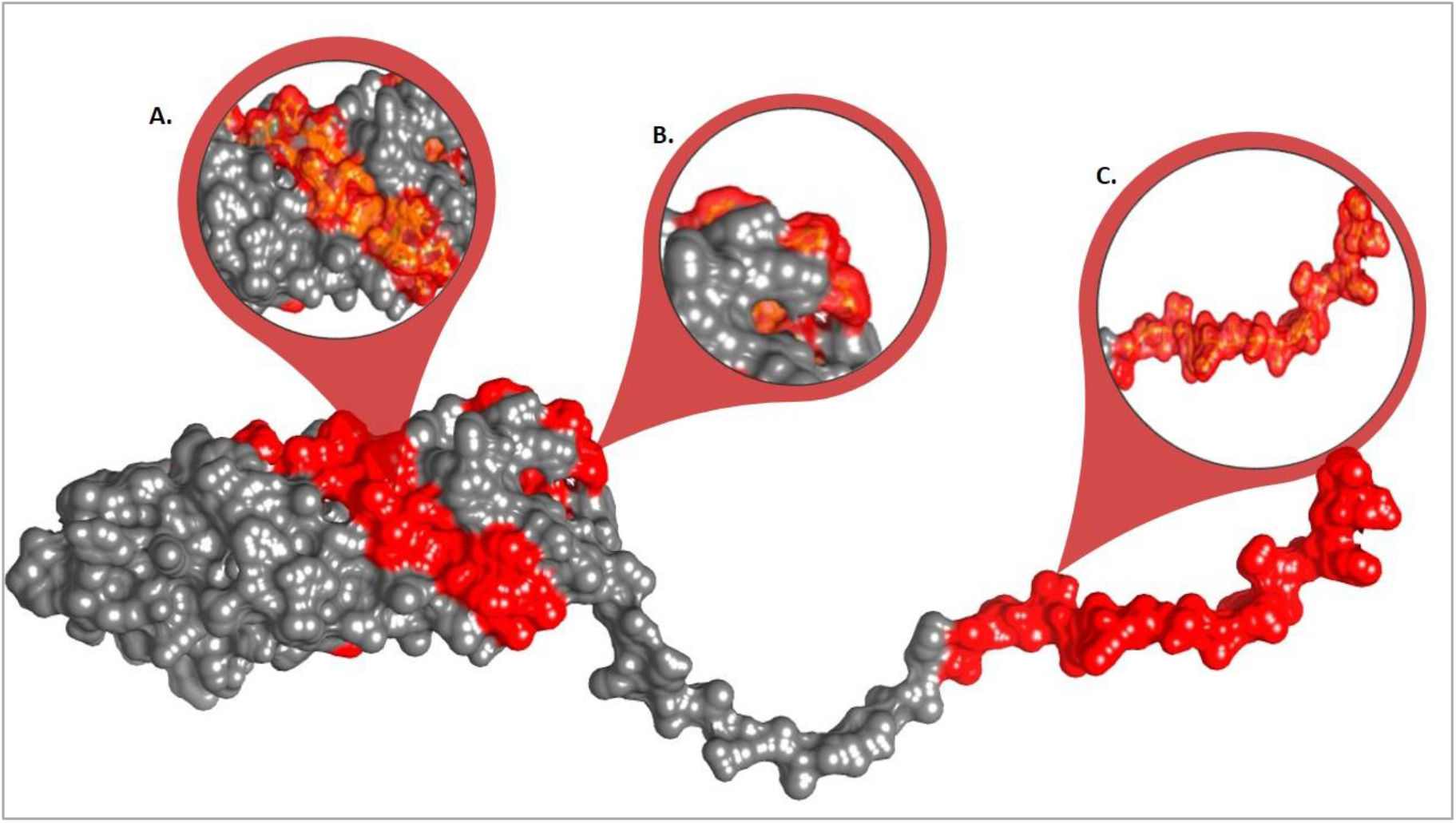
Three-Dimensional structure of OmpW protein of *B. abortus* showing most promising T- Cell peptides which interacts with MHC-II alleles, and located in the same conserved regions. A. shows peptides located in the region between positions 78 to 106. B. shows peptides located in the region between positions 169 to 183 and C. shows peptides located in the region between positions 1 to 21 (using chimera 1.13.1rc).

Many studies had predicted peptide vaccines for different microorganisms such as *madurella mycetomatis, HIV, pulmonary tuberculosis, treponema pallidum, pseudomona aeruginosa, COVID19* and *malaria* using immunoinformatics tools [22-28]. Many different approaches to reduce brucellosis risk were tested over the years like killed vaccines, subunit vaccines using recombinant proteins, or vector vaccines, with variable success [29]. Currently, vaccines for *Brucella* are made with live cells, so the use of OmpW family protein of *B. abortus* would be much safer [30]. In the present study, we identified several B and T cell epitopes that could be used as targets for vaccine design against *B. abortus*. However, several *in vivo* and *in vitro* studies will be needed in the future to confirm these results.

## 5. Conclusion

Vaccination is one of the most successful public health measures to reduce the burden of disease. Moreover, using *insilico* prediction methods will reduce the cost, time and effort needed to design these vaccines. In this work, we presented different peptides that can produce effective immunity in human against OmpW family protein of *B. abortus* for the first time. Seven B cell epitopes passed the antigenicity, accessibility, hydrophilicity and allergenicity tests. Seven promising MHC I epitopes were chosen with highest population coverage, while twenty four peptides for MHC II were chosen as the best targets for the vaccine. An astonishing and surprising combined population coverage for both MHC I and MHC II of 100% was obtained through this vaccine design.

## Data Availability

The data which support our findings in this study are available from the corresponding author upon reasonable request.

## Acknowledgment

Many thanks go to the National Center for Biotechnology Information (NCBI), BioEdit, Immune Epitope Database (IEDB) Analysis Resource, RaptorX server, and UCSF Chimera team.

## References

1. Wafa Al-Nassir, M.V.L., Robert A Salata, Nicholas John Bennett, Brucellosis Treatment & Management. 2018.

2. Al Dahouk, S., et al., Seroprevalence of brucellosis, tularemia, and yersiniosis in wild boars (Sus scrofa) from north-eastern Germany. Journal of Veterinary Medicine, Series B, 2005. 52(10): p. 444–455.

3. Pappas, G., et al., The new global map of human brucellosis. The Lancet infectious diseases, 2006. 6(2): p. 91–99.

4. Percin, D., Microbiology of Brucella. Recent Pat Antiinfect Drug Discov, 2013. 8(1): p. 13–7.

5. Golding, B., et al., Immunity and protection against Brucella abortus. Microbes and infection, 2001. 3(1): p. 43–48.

6. Gwida, M., et al., Brucellosis–regionally emerging zoonotic disease? Croatian medical journal, 2010. 51(4): p. 289–295.

7. Corbel, M.J., Brucellosis in humans and animals. 2006: World Health Organization.

8. McClean, S., Eight stranded β-barrel and related outer membrane proteins: role in bacterial pathogenesis. Protein and peptide letters, 2012. 19(10): p. 1013–1025.

9. Vemulapalli, T.H., et al., Role in virulence of a Brucella abortus protein exhibiting lectin-like activity. Infection and immunity, 2006. 74(1): p. 183–191.

10. Jezi, F.M., et al., Immunogenic and protective antigens of Brucella as vaccine candidates. Comparative Immunology, Microbiology and Infectious Diseases, 2019. 65: p. 29–36.

11. Jespersen, M.C., et al., BepiPred-2.0: improving sequence-based B-cell epitope prediction using conformational epitopes. 2017. 45(W1): p. W24–W29.

12. Kolaskar, A. and P.C.J.F.l. Tongaonkar, A semi-empirical method for prediction of antigenic determinants on protein antigens. 1990. 276(1-2): p. 172–174.

13. Emini, E.A., et al., Induction of hepatitis A virus-neutralizing antibody by a virus-specific synthetic peptide. 1985. 55(3): p. 836–839.

14. Parker, J., D. Guo, and R.J.B. Hodges, New hydrophilicity scale derived from high-performance liquid chromatography peptide retention data: correlation of predicted surface residues with antigenicity and X-ray-derived accessible sites. 1986. 25(19): p. 5425–5432.

15. Nielsen, M., et al., Reliable prediction of T- Cell epitopes using neural networks with novel sequence representations. 2003. 12(5): p. 1007–1017.

16. Andreatta, M. and M.J.B. Nielsen, Gapped sequence alignment using artificial neural networks: application to the MHC class I system. 2015. 32(4): p. 511–517.

17. Nielsen, M. and O.J.B.b. Lund, NN-align. An artificial neural network-based alignment algorithm for MHC class II peptide binding prediction. 2009. 10(1): p. 296.

18. Dimitrov, I., D.R. Flower, and I. Doytchinova. AllerTOP-a server for in silico prediction of allergens. in BMC bioinformatics. 2013. BioMed Central.

19. Bui, H.-H., et al., Predicting population coverage of T- Cell epitope-based diagnostics and vaccines. 2006. 7(1): p. 153.

20. Kazi, A., et al., Current progress of immunoinformatics approach harnessed for cellular-and antibody-dependent vaccine design. Pathog Glob Health, 2018. 112(3): p. 123–131.

21. Desai, D.V. and U. Kulkarni-Kale, T- Cell epitope prediction methods: an overview. Methods Mol Biol, 2014. 1184: p. 333–64.

22. A Multiple Peptides Vaccine against COVID-19 Designed from the Nucleocapsid phosphoprotein (N) and Spike Glycoprotein (S) via the Immunoinformatics Approach. Informatics in Medicine Unlocked, 2020: p. 100476.

23. Elhag, M., et al., Design of Epitope-Based Peptide Vaccine against Pseudomonas aeruginosa Fructose Bisphosphate Aldolase Protein Using Immunoinformatics. Journal of Immunology Research, 2020. 2020: p. 9475058.

24. Mohammed, A.A., et al., Epitope-Based Peptide Vaccine Against Fructose-Bisphosphate Aldolase of Madurella mycetomatis Using Immunoinformatics Approaches. Bioinform Biol Insights, 2018. 12: p. 1177932218809703.

25. Chatterjee, N., et al., Scrutinizing Mycobacterium tuberculosis membrane and secretory proteins to formulate multiepitope subunit vaccine against pulmonary tuberculosis by utilizing immunoinformatic approaches. Int J Biol Macromol, 2018. 118(Pt A): p. 180–188.

26. Cravo, P., et al., In silico epitope mapping and experimental evaluation of the Merozoite Adhesive Erythrocytic Binding Protein (MAEBL) as a malaria vaccine candidate. Malar J, 2018. 17(1): p. 20.

27. Pandey, R.K., et al., Immunoinformatics approaches to design a novel multi-epitope subunit vaccine against HIV infection. Vaccine, 2018. 36(17): p. 2262–2272.

28. Elhag, M., et al., Immunoinformatics Approach for Designing an Epitope-Based Peptide Vaccine against Treponema pallidum Outer Membrane Beta-Barrel Protein. Immunome Research, 2020. 16(2): p. 1–12.

29. Kim, W.K., et al., Protective efficacy of an inactivated Brucella abortus vaccine candidate lysed by GI24 against brucellosis in Korean black goats. Can J Vet Res, 2019. 83(1): p. 68–74.

30. Araiza-Villanueva, M., et al., Proteomic Analysis of Membrane Blebs of Brucella abortus 2308 and RB51 and Their Evaluation as an Acellular Vaccine. Front Microbiol, 2019. 10: p. 2714.

